# Assessing PARP trapping dynamics in ovarian cancer using a CRISPR-engineered FRET biosensor

**DOI:** 10.1101/2025.03.12.642798

**Authors:** Dan Marks, Edwin Garcia, Sunil Kumar, Katie Tyson, Caroline Koch, Aleksander Ivanov, Joshua Edel, Hasan B. Mirza, William Flanagan, Christopher Dunsby, Paul M.W French, Iain A. McNeish

## Abstract

Poly(ADP-ribose) polymerase inhibitors (PARPi) have revolutionised the treatment of ovarian high grade serous carcinoma (HGSC), especially those with defective homologous recombination. However, the emergence of resistance poses a critical challenge, as over 50% of patients relapse within three years. The mechanisms underlying changes in PARP trapping, a central aspect of PARPi efficacy, are not well understood due to limitations in current experimental methodologies. Existing techniques lack resolution and throughput, impeding efforts to study PARP trapping dynamics over time with single-cell resolution. Effective tools to study PARP trapping in live cells are urgently needed to elucidate resistance mechanisms and inform therapeutic strategies.

To address this, we used CRISPR-Cas9 gene editing to dual-label endogenous *PARP1* with EGFP and mCherryFP in OVCAR4 cells to develop a novel intramolecular FRET-based biosensor that enables real-time, single-cell visualization of PARP trapping dynamics in live cells. High-content fluorescence lifetime imaging microscopy (FLIM) revealed dose dependent PARP trapping upon exposure to PARP inhibitor and distinguished between the trapping efficiencies of four different PARPi (veliparib, olaparib, rucaparib, talazoparib). Moreover, we found reduced PARP trapping in PARPi-resistant models, both *in vitro* and *in vivo*, providing critical evidence for altered PARP trapping as a resistance mechanism and illustrating the potential of this FRET biosensor to interrogate resistance mechanisms quantitatively.

This PARP trapping biosensor represents a transformative advance, enabling dynamic, high-resolution analysis of mechanisms underlying cancer drug resistance. It provides critical insights into the heterogeneity of PARPi resistance, with implications for developing more effective therapies and improving personalised treatment strategies for ovarian cancer patients.

**Graphical abstract:** 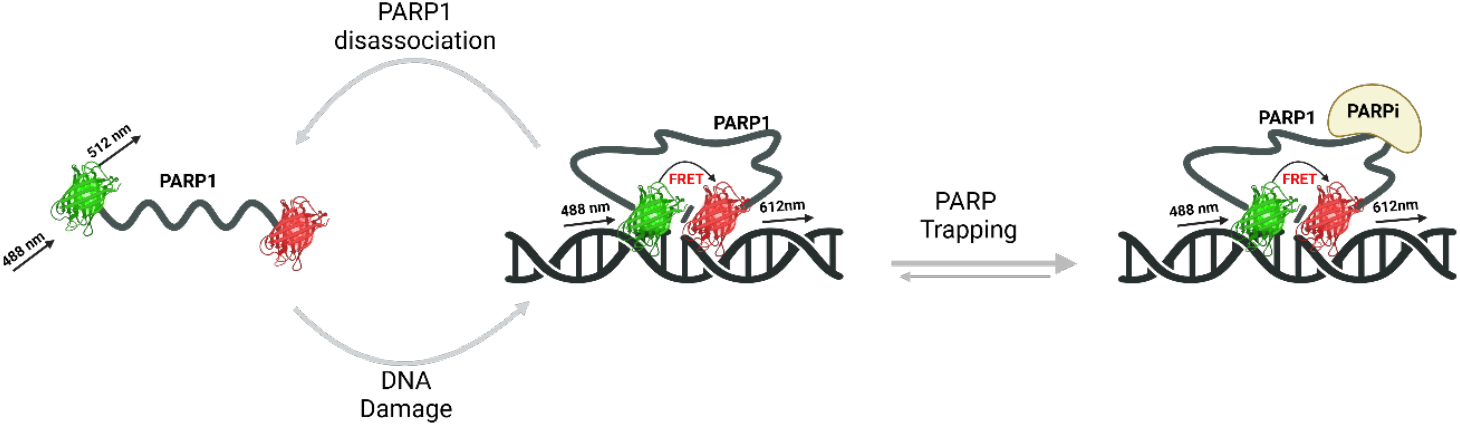

## Introduction

High-grade serous carcinoma (HGSC) is the most common and lethal subtype of ovarian cancer. Poly(ADP-ribose) polymerase inhibitors (PARPi) have transformed the management of *BRCA1/2*-mutated and homologous recombination (HR) defective high-grade ovarian carcinoma^1,2^. They also have activity in HR proficient disease^3,4^. However, more than 50% of patients relapse within three years of treatment initiation, highlighting the need to understand and overcome PARPi resistance.

Upon formation of single-strand DNA breaks, PARP1 attaches to damage lesions within seconds. NAD^+^ is used as a substrate to form PAR chain modifications (auto-PARylation), leading to chromatin remodelling and the recruitment of repair factors. PARPi function by preventing auto-PARylation, causing PARP to become trapped on the DNA, which increases the steric complexity of lesion sites, resulting in the formation of double-strand DNA breaks (DSBs) during replication. When HR is functional, these DSBs can be repaired accurately. However, in cells lacking functional HR, such as those with *BRCA1/2* mutations, this can lead to genomic instability and cell death over multiple cell cycles^5,6^.

Multiple mechanisms of PARPi resistance have been identified in preclinical models, although few have been validated in clinical settings^7^. Reactivation of functional HR, often due to *BRCA1/2* reversion mutations, has been identified as a key mechanism of PARPi resistance^8^. Other proposed mechanisms include increased drug efflux (MDR1-dependent)^9^, downregulation of poly(ADP-ribose) glycohydrolase (PARG)^10^, loss of 53BP1 expression^11,12^ and diminished PARP trapping due to mutations in *PARP1*^13^.

Current methodologies to assess PARP trapping include chromatin immunoprecipitation (ChIP)^14^, which requires fixed, pooled cell populations, limiting its applicability to dynamic analyses. While fluorescence recovery after photobleaching (FRAP) partially overcomes this by enabling measurements in live cells, it is restricted to small populations (<100 cells), lacks the throughput needed to capture heterogeneity in response and typically relies on overexpression of fluorescently tagged PARP1, which can alter protein kinetics.

Here, we address these limitations by developing a novel intramolecular FRET biosensor for PARP trapping by labelling endogenous PARP1 with both EGFP and mCherryFP, enabling real-time, single-cell resolved measurements of PARP trapping via high-content fluorescence lifetime imaging microscopy (FLIM). We demonstrate the utility of this biosensor by showing reduced PARP trapping in both *in vitro* and *in vivo* models of PARPi resistance. These findings pave the way for a deeper understanding of resistance mechanisms and inform the development of more effective and durable therapies for ovarian cancer patients.

## Methods

### Cell Culture

OVCAR4 cells were obtained from ATCC and were cultured in DMEM (Gibco 11965092), supplemented with 10% heat-inactivated foetal bovine serum (Gibco 10270-106), 2 mM L-Glutamine (Thermo Fisher Scientific 25030-081) and 100 ug/mL penicillin/streptomycin (P/S) (Thermo Fisher Scientific 15070-063). All cells were grown in 5% CO_2_, 37°C with humidity, and used for a maximum of 10 passages. Cell identity was validated by 16 locus STR validation and tested for mycoplasma infection regularly using MycoAlert™ detection kit (Lonza LT07-218).

### CRISPR/Cas9 knock-in of EGFP and mCherryFP at endogenous *PARP1* locus

Open-access python package pyDNA was used for in silico plasmid sequence design. The software program CHOPCHOP (https://chopchop.rc.fas.harvard.edu/) was used to design guide RNA (gRNA) targeted to the N- or C-termini of *PARP1*. 5’ and 3’ homology arms (HA) to *PARP1* were synthesised by PCR.

For C-terminal labelling, the mCherry fragment was amplified from a mCherry-H2B plasmid (Addgene 20972) with complementary overhangs to the 5’ and 3’ HA. The targeting vector (kindly gifted by Matsuda et al^15^) backbone was digested with AscI (Thermo Fisher Scientific ER1891) and SwaI (Thermo Fisher Scientific ER1241), isolating the 2004 bp backbone fragment. The 5’ and 3’ homology arms (sequences taken from Perez-Leal et al. ^16^) were amplified via PCR, using primers designed to create fragments with complementary overhangs ∼22 bp in length. Gibson assembly reactions were performed at 50 °C for 1 h using the NEBuilder HiFi DNA assembly cloning kit (New England Biolabs E5520S), combining DNA fragments with overlapping sequences at equal molar ratios with the digested backbone sequence. The reaction product transformed in NEB 5α competent *E. coli* (New England Biolabs C2987) and isolated using QIAprep Spin Miniprep Kit (Qiagen 27104). This PARP1 C-terminal targeting vector was used with the same gRNA previously used by Perez-Leal et al., GTCAATTTTAAGACCTCCCTG, which was ligated into pX330-U6-chimeric-BB-Cbh-hSpCas9 plasmid (Addgene 42230)^16^.

For N-terminal knock-in (KI), *in trans* paired nicking was used. Two plasmids kindly gifted by Chen et al^17^ (pgRNA^PAPR1^, pDonor^PARP1^) were subsequently transfected with pST1374-N-NLS-flag-linker-Cas9-D10A (Addgene 51130). These three plasmids constituted the CRISPR system for N-terminal targeting of PARP1.

For CRISPR transfections, 2×10^5^ OVCAR4 cells were plated overnight in 6-well plates in antibiotic-free medium and transfected with 1 μg of each plasmid using Viafect (Promega E4981). Fluorescence positive cells were isolated after 24–48h using FACS, and single cell colonies were expanded.

### RT-qPCR

Culture medium was aspirated from 6-well plates and 350 μL RLT buffer was added and frozen at -80°C. Plates were thawed on ice and 70% ethanol was added and transferred into a RNeasy Micro Kit column (Qiagen, 74104). RNA extraction was performed as per the manufacturer’s instructions, genomic DNA was digested using RNase-Free DNase (Qiagen, 79254). RNA was eluted in 30 μL nuclease-free water, concentration and quality were estimated using Nanodrop and Qubit 3.0 analysis.

1 μg RNA was used in each 20 μL cDNA reaction with the High-Capacity cDNA Reverse Transcription Kit (Applied Biosystems, 4368814) with cycles of 25°C 10 minutes, 37°C 120 minutes, and 85°C 5 minutes. The resulting cDNA was diluted in 140 μL nuclease-free water. qRT-PCR reaction was performed using 9 μL cDNA, 1 μL primer and 10 μL TaqMan Universal Master Mix II no UNG (Thermo Fisher Scientific, 4440040). TaqMan primer probes (*GAPDH* (Hs02786624_g1), *PARG (*Hs00608254_m1*), TP53BP1* (Hs00996827_m1), *ABCB1* (Hs00184500_m1), Thermo Fisher Scientific). Samples were loaded in a 96-well plate (Applied Biosystems, 4311971) and sealed with optical plate seal (Applied Biosystems, 4346907) and analysed on a StepOnePlus (Applied Biosystems). Gene expression was normalised to *GAPDH*

### Long-read direct RNA-sequencing

For Oxford Nanopore Technologies (ONT) direct RNA sequencing (DRS), 1 μg total RNA was used to generate libraries using the Direct RNA Sequencing Kit (Oxford Nanopore Technologies, SQK-RNA004) according to the manufacturer’s instructions. Sequencing was performed using a MinION device (ONT, MIN-101B) with RNA flow cells (ONT, FLO-MIN004RA). Base calling was performed with MinKNOW using Dorado. Basecalled sequencing data were processed using the wf-transcriptomes pipeline within the Epi2Me software. The workflow includes read preprocessing, alignment using minimap2, transcript assembly and quantification, quality control and validation. Reads were visualised using Integrative Genomics Viewer (IGV, www.igv.org) and a ratio of reads per isoform of PARP1 was calculated per sample.

### Automated wide-field time-gated FLIM microscope and FLIM data acquisition

The instrument used for these experiments is shown in **Figure 2**, and described in detail in ^18^. Quasi-confocal FLIM was implemented upon a motorized inverted epifluorescence microscope (Olympus IX-81, incorporating a ZDC autofocus) with a spinning disc scanner (Yokogawa, CSU-X) used for all experiments. Upon drug treatment, imaging 96-well plates were immediately transferred to the microscope chamber at 37 °C, 5% CO_2_, mounted on a motorized x-y stage (Märzhäuser Wetzlar GmbH) and left for 1 h for temperature equalisation before optically sectioned FLIM acquisitions. A 20x air objective lens (Olympus UPlanApo 20x) with an NA of 0.7 was used. For excitation of EGFP at 488 nm, the output beam from a tuneable femtosecond Ti:Sapphire laser (Mai Tai HP, Spectra-Physics) was directed to a second harmonic generation unit (Spectra-Physics, 3980) to provide pulsed excitation radiation at a 80 MHz repetition rate.

1×10^4^ FRET biosensor-expressing OVCAR4 cells were seeded per well onto a fibronectin (Sigma F0895) coated (3 μg/cm^2^) 96-well glass imaging plate (Greiner 655892) in CO_2_-dependent imaging medium 48 h before imaging and maintained at 37°C with 5% CO_2_ during imaging. A 488/532 dichroic mirror (Semrock, Di01-T488/532-13×15×0.5) directed the excitation beam, with power adjusted to ∼200 μW at the sample plane. Fluorescence emission was collected through a 520/35 nm filter (Semrock, FF01-520/35-25) and detected using a gated optical intensifier (Kentech Instruments, HRI-HL) coupled to a cooled CCD camera (Photometrics Retiga R1) via a 0.7× demagnification relay. The gating signal was set to a 4 ns width and synchronized to the laser pulses, with time delays controlled by the μManager plugin openFLIM-HCA (https://github.com/imperial-photonics/openFLIM-HCA). FLIM images were acquired at seven time delays with a total FLIM data acquisition time of ∼10 s per field of view (FOV).

Data were saved as OME-TIFF files and analysed using the open-source MATLAB software *FLIMfit*^*19*^. The instrument response function (IRF) was accounted for in each experiment by measuring the fluorescence decay of a reference dye (75 μM Coumarin 6 in ethanol) at 25 ps intervals over a 12.5 ns pulse period.

Additionally, quenched fluorescein (2.5 μL of 250 μM fluorescein in 100 μL 2M potassium iodide) was used as a secondary reference with a ∼100 ps lifetime. The Coumarin 6 image dataset was fitted using reference reconvolution employing the spatially averaged quenched fluorescein decay as the reference decay. The fluorescence lifetime of the quenched fluorescein was input as 180 ps as determined from a previous time-correlated single-photon counting measurement. The FLIMfit option “Create IRF Shift Map” was used to calculate the spatially varying IRF. This shift map was used subsequently when fitting FLIM data to correct for spatial variations in the IRF across the HRI FOV. Measurements were taken before and after each imaging session to confirm stability. A time-varying background was measured in well containing only cell culture media. Global double-exponential fitting was performed per field of view (FOV), and weighted mean lifetime values were extracted on a per-cell, per-condition basis.

### Imaging intrinsic fluorescence of rucaparib

A custom-built multi-beam, multi-photon, multi-well plate microscope (M^3^M) built on the opensource *openFrame* microscope (Cairn Research, UK) was used. Prior to imaging, cells were treated for 1 h with 10 μM rucaparib, which was excited at 365 nm to image intracellular rucaparib using a 20X 0.95NA water immersion objective (Nikon, MRD77200). Scripts were written to use CellPose Cyto3 model for automated whole cell segmentation in Fiji to define ROIs within which the rucaparib total intensity was quantified to assess single cell intracellular drug concentration.

### *In vivo* experiments

*In vivo* experiments were performed at the Central Biological Services facility, Imperial College London and approved by the Imperial College Animal Welfare & Ethics Review Body (AWERB). Experiments were performed under the authority of project licence numbers PA780D61A and PP1321516. All experiments conformed to UK Home Office regulations under the Animals (Scientific Procedures) Act 1986, including Amendment Regulations 2012.

5×10^6^ OVCAR4 GFP-PARP1-mCherryFP cells were injected intraperitoneally (I.P.) in 200 μL PBS into 6-7 week old female CD1-nude (Crl:CD1-*Foxn1*^*nu*^) mice (Charles River, U.K.). On days 15–28 inclusive, 25 mg/kg olaparib (Selleckchem S1060) dissolved in 18% SBE-β-CD (MedChemExpress HY-17031), 10% DMSO in saline was injected I.P. Mice were culled when the first in each group reached humane endpoint and tumours harvested into PBS on ice. If there was no ascites present, peritoneal lavage (5 mL of 2 mM EDTA in PBS) was performed to isolate ascites spheroids, which were centrifuged at 200 x g for 5 min and resuspended in TrypLE Express Enzyme (Gibco 12604013) for 10 min, 37°C. Disassociated spheroids were subsequently resuspended in medium and cultured under standard conditions.

### Statistical analyses

All statistical tests were performed using Prism v.9.4.1 (GraphPad). A p value ≤0.05 was considered statistically significant.

### Data and materials availability

All raw data and code are available upon request. Ethics statement

All *in vivo* experiments performed in mice were approved by the Animal Welfare & Ethics Review Body (AWERB) at Imperial College London. Experiments were performed under the project license numbers PA780D61A and PP1321516 at Imperial College London. All experiments conformed to UK Home Office regulations under the Animals (Scientific Procedures) Act 1986, including Amendment Regulations 2012.

## Results

### Endogenous dual-labelling of PARP1 using CRISPR-Cas9

To generate a dual-labelled PARP1 fluorescence resonance energy transfer (FRET) biosensor, a sequential CRISPR knock-in (KI) strategy was employed, targeting the N- and C-termini of the *PARP1* locus, based on prior studies^20^ (**Figure 1A**). The initial CRISPR KI utilised a system provided by Chen et al^17^, comprising two plasmids (pDonor^PARP1.TS^ and SpCas9^D10A^:gRNA^PARP1^) for *in trans* paired nicking to insert EGFP at the N-terminus of PARP1. OVCAR4 cells were transfected with these plasmids, generating a heterozygous population of EGFP-PARP1 and wildtype (WT) cells. Fluorescence-activated cell sorting (FACS) was used to isolate EGFP-positive cells, yielding a homogenous OVCAR4 EGFP-PARP1 labelled cell line.

**Figure 1:**
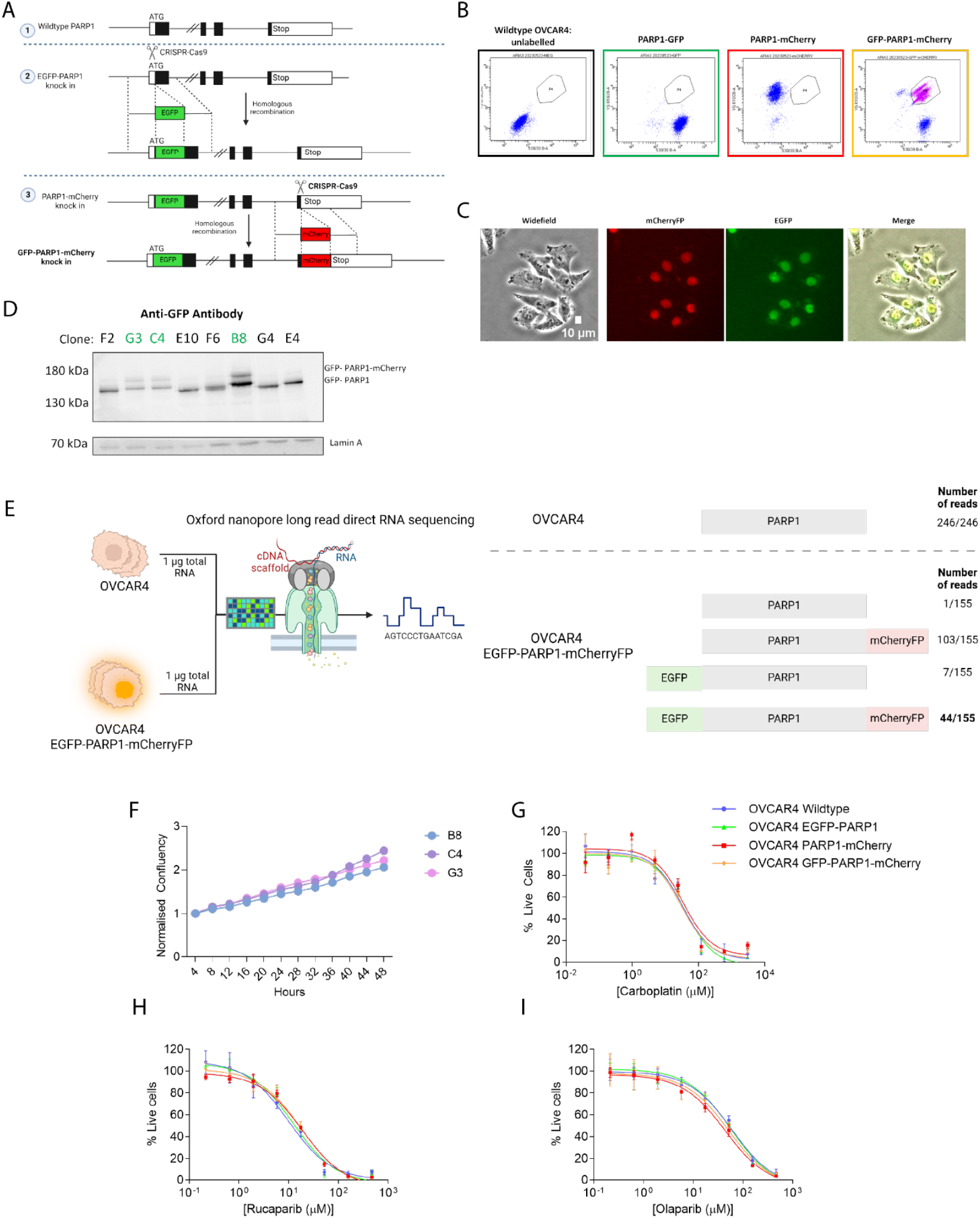
Endogenous dual-labelling of PARP1 with donor (EGFP) and acceptor (mCherryFP) pair using CRISPR-Cas9. **A:** Schematic of the sequential gene editing approach to knock in EGFP at the N-terminal of PARP1, followed by mCherry at the C-terminal. **B:** Dual-labelled OVCAR4 cells were isolated by FACS. **C:** Representative images of dual-labelled OVCAR4 EGFP-PARP1-mCherryFP cells showing phase, mCherryFP, EGFP and merged images, scale bar = 10 μm. **D:** Long-read direct RNA sequencing was achieved using oxford nanopore sequencing. RNA was extracted from parental OVCAR4 WT cells and CRISPR edited EGFP-PARP1-mCherryFP cells. The isoform frequencies for each population are shown below. **E:** Immunoblots were performed to assess labelling of PARP1 protein; clones highlighted in green (G3, E4 and B8) showed dual labelled EGFP-PARP1-mCherryFP. **F**: Growth rate of each clone was assessed by acquiring timelapse images over 48 h using the Incucyte S3 system at 10X magnification. Segmentation and confluency analysis performed in Incucyte software. Data were normalised to the confluency at t=0 for each subsequent time point. **G-I:** SRB assays for OVCAR4 WT compared to the 3 CRISPR generated cell lines treated with olaparib (G), carboplatin (H), or rucaparib (I). OVCAR4 wild-type (blue), OVCAR4 EGFP-PARP1 (green), OVCAR4 PARP1-mCherryFP (red), OVCAR4 EGFP-PARP1-mCherryFP (orange). Values were normalised to untreated controls to calculate live cell values per treatment. One-way ANOVA was performed with multiple comparisons comparing the viability of each line to the parental cell line, all were non-significantly different, n=3.

A second round of CRISPR KI targeted the C-terminus of *PARP1* to insert mCherryFP. A rhodopsin targeting vector (TV)^15^ was modified via restriction digestion and Gibson assembly, replacing the original EGFP sequence with mCherryFP and adapting the homology arms for the *PARP1* locus. The modified TV was co-transfected with gRNA enabling C-terminal PARP1-mCherryFP labelling. FACS was again performed to isolate cells co-expressing EGFP and mCherryFP (**Figure 1B,C**). From the heterogeneous dual-labelled population, single-cell clones were isolated, and western blot analyses confirmed that three clones expressed both EGFP and mCherryFP (170 kDa band, **Figure 1D**).

To confirm knock-in, long-read direct sequencing was performed (Oxford Nanopore Technologies (ONT)) using RNA extracted from parental OVCAR4 and OVCAR4 EGFP-PARP1-mCherryFP cells (clone B8). This showed a single PARP1 isoform in the wild-type population (246/246 reads). In the CRISPR-edited cells (**Figure 1E**) PARP1-mCherryFP was identified in 103/155 reads, EGFP-PARP1 in 7/155 reads and EGFP-PARP1-mCherryFP in 44/155 reads. This suggests that the dual-labelled PARP1 constitutes at least ∼30% of the total *PARP1* transcript pool. Dual labelling neither altered morphology or cellular growth rates (**Figure 1C,F**), nor sensitivity to carboplatin (**Figure 1G**), or PARP inhibitors rucaparib or olaparib (**Figure 1H,I**), validating the use of these models for subsequent studies.

### FLIM-FRET Analysis Validates Dual-Labelled OVCAR4 Cells as a Biosensor for PARP Trapping

To evaluate the efficacy of dual-labelled OVCAR4 EGFP-PARP1-mCherryFP cells as a FRET biosensor, wide-field time-gated fluorescence lifetime imaging microscopy (FLIM) high-content analysis (HCA) was performed (**Figure 2A**). Fluorescence lifetime images of EGFP (the FRET donor) were globally fitted using FLIMfit software^19^. Regions of interest (ROIs) corresponding to individual nuclei were segmented to enable single-cell quantification and population-level analysis under different treatment conditions. An initial experiment assessed the response to changes in DNA damage and PARP trapping following treatment with carboplatin and rucaparib, respectively (**Figure 2B,C**). Fluorescence lifetime reductions were analysed at both the field-of-view (FOV) and single-cell levels, revealing consistent trends. Carboplatin treatment induced non-significant reductions in mean fluorescence lifetime compared to untreated cells. This was not unexpected, as platinum drugs induce DNA crosslinking, leading to dsDNA breaks, which are not the target of PARP1-mediated repair. In contrast, rucaparib caused significant dose-dependent reductions in fluorescence lifetime, with mean values of 2532 ps and 2516 ps at 10 μM and 30 μM rucaparib, respectively (*p* < 0.05). This suggests that the FRET biosensor reports changes in PARP trapping specifically, rather than DNA damage generally. Notably, FOV- and single-cell analyses were highly consistent, with only a 0.07% average variation, confirming the reliability of population-level fluorescence lifetime measurements. γH2AX phosphorylation immunofluorescence showed a dose dependent increase in γH2AX foci per cell, confirming that 1 h exposure to PARPi was sufficient to induce DNA damage (**Figure 2D**).

**Figure 2:**
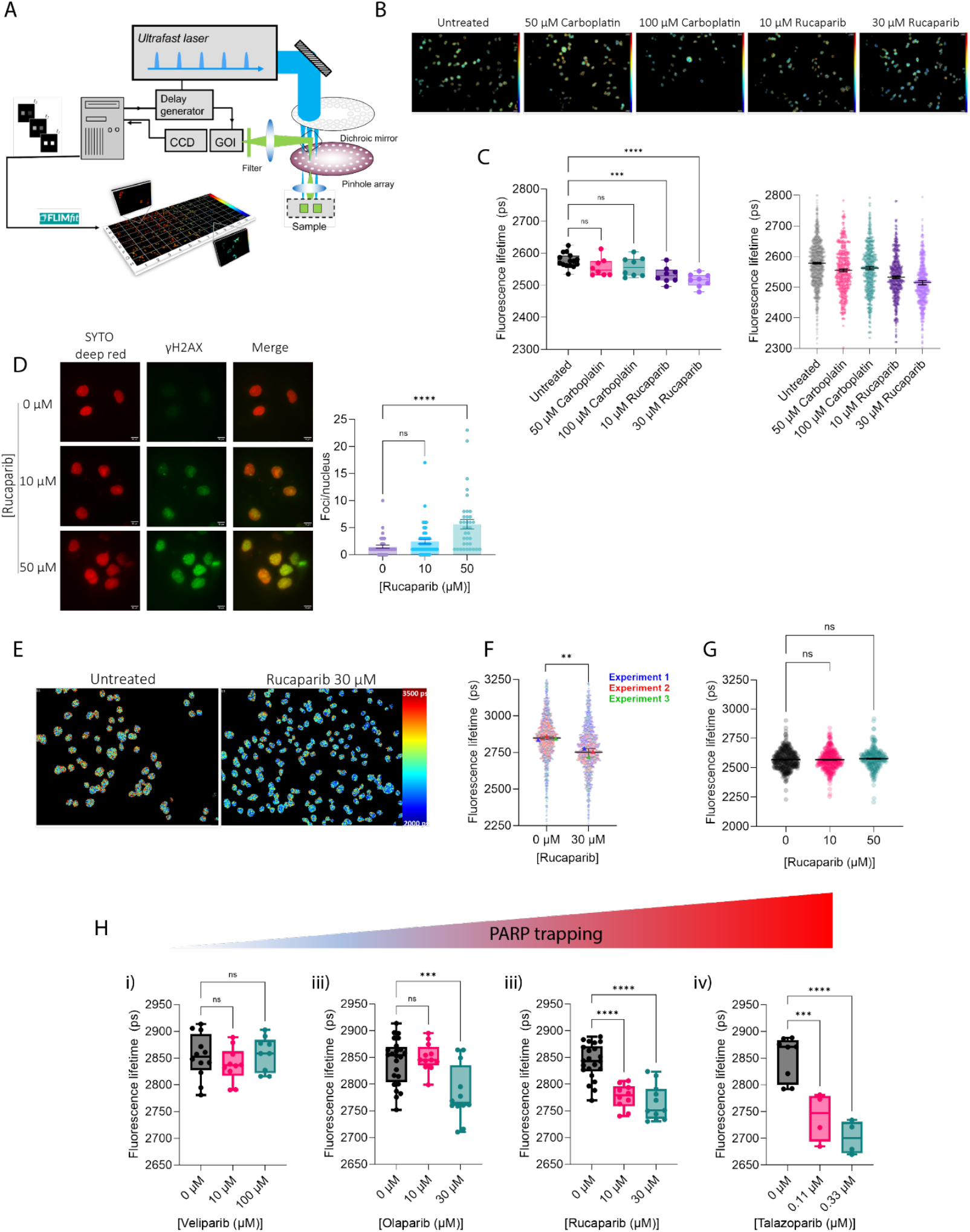
EGFP-PARP1-FRET biosensor detects changes in PARP trapping, rather than DNA damage. **A:** System diagram of wide-field time-gated fluorescence lifetime imaging microscope. Intensity images are acquired at various time-gates and fluorescence decays are fitted using global fitting of a double-exponential decay in FLIMfit. **B:** Representative FLIM images per condition from HCA FLIM-FRET assay, images are pseudo-coloured by mean fluorescence lifetime. **C:** Fluorescence lifetime of FRET biosensor donor upon treatment with carboplatin (50, 100 μM) or rucaparib (10, 30 μM) for 1 h, compared to untreated controls. Values are summarised as FOV averages (left) or single-cell mean fluorescence lifetimes (right). One-way ANOVA (Kruskal-Wallis) with Dunn’s multiple comparison test was used to assess statistical significance between treatment conditions ***; p<0.0001, ****; p<0.0001. **D:** Representative images of γH2AX foci per cell following 1 h of rucaparib exposure at 0, 10 or 50 μM in OVCAR4 EGFP-PARP1-mCherryFP cells, quantification shown with mean and SEM values. Scale bar=10 μm, N=3 biological replicates. **E:** Representative images of segmented OVCAR4 EGFP-PARP1-mCherryFP nuclei, pseudo-coloured by mean fluorescence lifetime values with untreated (left) and treated with rucaparib 30 μM rucaparib for 1 h values shown in **F. G:** Single-cell mean fluorescence lifetime values are plotted per condition and colour-coded by the experimental repeat they are part of. Fluorescence lifetime of EGFP in donor only controls (OVCAR4 EGFP-PARP1) exposed to 0, 10 or 50 μM rucaparib for 1 h. Statistical significance between treatment conditions were assessed using an unpaired t-test ns; p>0.05, **; p<0.01, n=3 biological repeats. **H:** Change in FRET biosensor donor fluorescence lifetime in OVCAR4 EGFP-PARP1-mCherryFP expressing live cells, upon treatment for 1 h with differing PARPi, veliparib (i), olaparib (ii), rucaparib (iii) or talazoparib (iv) at various doses. One-way ANOVA (Kruskal-Wallis) with Dunn’s multiple comparison test was used to assess statistical significance between treatment conditions.

Separate repeat measurements carried out at weekly intervals corroborated the reproducibility of changes in PARP trapping upon rucaparib treatment (**Figure 2E,F**). Statistically significant reductions in fluorescence lifetime were consistently observed in biosensor-expressing cells treated with 30 μM rucaparib for 1 h compared to untreated controls. Conversely, OVCAR4 EGFP-PARP1 (donor only) cells did not exhibit significant fluorescence lifetime changes under identical conditions, underscoring the specificity of the biosensor (**Figure 2G**).

To evaluate the biosensor’s capacity to differentiate the trapping potency of different PARPi, cells were treated with veliparib, olaparib, rucaparib or talazoparib. Distinct fluorescence lifetime profiles were observed, consistent with published rankings of PARP trapping efficiency (**Figure 2H**). Veliparib resulted in no significant difference in mean lifetime values (2854, 2838 and 2858 ps for 0, 10 and 100 μM respectively), indicating minimal PARP trapping. Olaparib and rucaparib both induced dose-dependent reductions in fluorescence lifetime (Olaparib: 2842, 2850, 2782 and 2762 ps for 0, 10, 30 and 50 μM, respectively; rucaparib: 2842, 2777, 2766, and 2765 ps for 0, 10, 30 and 50 μM, respectively). The reductions were statistically significant at 30 μM (olaparib) and ≥10 μM (rucaparib). Talazoparib exhibited the most potent PARP trapping effects, with significant reductions in fluorescence lifetime detected at 0.11 μM (2850, 2740 and 2701 ps for 0, 0.11, 0.33 μM, respectively; p< 0.01). These findings validate that dual-labelled PARP1 FRET can discern PARP trapping in live cells with high sensitivity and reproducibility. The rank order of PARPi trapping potency observed here, talazoparib > rucaparib ≈ olaparib > veliparib, closely aligns with prior literature^14^, further supporting the biosensor’s utility for investigating PARP-trapping in live cells.

### Decreased sensitivity to PARPi correlates with lower PARP trapping *in vitro*

To induced resistance *in vitro*, OVCAR4 EGFP-PARP1-mCherryFP cells were treated daily with 20 μM olaparib or rucaparib (doses approximating their 72h IC50 values) for nine weeks, with daily medium replenishment (**Figure 3A**). Populations derived from this protocol are termed “olaparib-exposed” and “rucaparib-exposed” in subsequent analyses. The sensitivity of these cells to their respective PARPi (**Figure 3B,C**) decreased significantly, with an increase in IC50 values from 22 μM to 36 μM (rucaparib) and from 23 μM to 39 μM (olaparib), confirming that the long-term treatment protocol effectively induced PARPi resistance. Both rucaparib- and olaparib-exposed cells also demonstrated reduced sensitivity to cisplatin (Error! Reference source not found.) the cisplatin IC50 in rucaparib-exposed cells was 17 μM, and 13 μM in olaparib-exposed cells, compared to 5 μM for parental cells.

**Figure 3:**
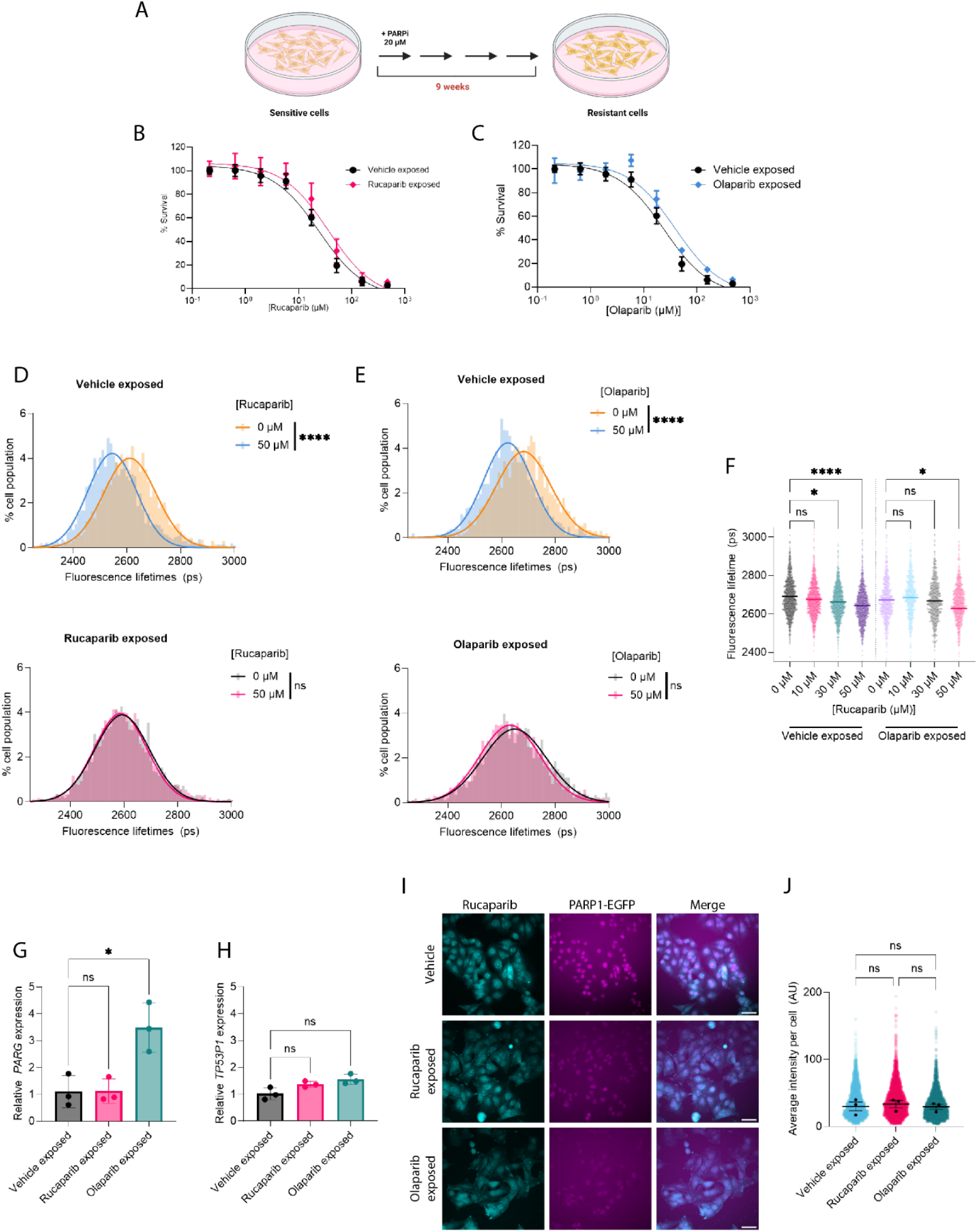
Induction of resistance in vitro induces a reduction in PARP trapping as assessed by EGFP-PARP1-mCherryFP FRET biosensor. **A:** Schematic of the treatment protocol to induce in vitro resistance to olaparib or rucaparib. OVCAR4 EGFP-PARP1-mCherryFP cells were treated daily with 20 μM olaparib or rucaparib for 8 weeks. Resistance was assessed by SRB assay to (**B**) rucaparib or (**C**) olaparib and compared to vehicle treated parental OVCAR4 EGFP-PARP1-mCherryFP cells. **D:** Live-cell FLIM-FRET HCA assays were performed to assess changes in donor lifetime of the EGFP-PARP1-mCherryFP FRET biosensor upon 1 h treatment with rucaparib or **E:** olaparib in sensitive and resistant populations. Linear regression (gaussian) curves were fitted to the single-cell FLIM data to show population level distributions of mean weighted fluorescence lifetime. **F:** Change in FRET biosensor donor fluorescence lifetime of olaparib sensitive (left) or resistant (right) cells upon treatment with rucaparib at 0, 10, 30 or 50 μM for 1 h. One-way ANOVA test was used to assess statistical significance between treatment conditions. **G:** RT-qPCR analysis of PARG and **H**: TP53BP1 relative mRNA expression normalised to GAPDH expression. N=3 biological repeats. **I:** Representative images of rucaparib and EGFP fluorescence in OVCAR4 EGFP-PARP1-mCherryFP cells following vehicle, rucaparib or olaparib exposure in vitro for 9 weeks. **J:** Quantification of average intracellular rucaparib intensity of images in **I**, each point is a single cell average and error bars represent population median. N=3 biological repeats.

To assess changes in PARP trapping, FLIM-FRET HCA assays were performed on PARPi-exposed cells. Following re-treatment with rucaparib, there was no significant change in fluorescence lifetime in the rucaparib-exposed population (2594 ps to 2589 ps, *p* = 0.9822), in contrast to a significant reduction in parental cells (2616 ps to 2553 ps, *p*<0.0001) (**Figure 3D**). A similar trend was observed in the olaparib-exposed population, where re-exposure to olaparib produced no significant change in fluorescence lifetime (2665 ps to 2656 ps, *p*<0.6836) whilst the parental population underwent a significant reduction in lifetime (2700 ps to 2664 ps, *p*<0.0001) (**Figure 3E**). To evaluate cross-resistance, olaparib-exposed cells were treated with rucaparib, which produced no statistically significant reduction in fluorescence lifetime at 50 μM (2672 ps and 2629 ps for 0 and 50 μM respectively (*p=0*.*111*). As previously, fluorescence lifetimes were significantly reduced in parental cells at 30 μM (2690 to 2644 ps; *p<0*.*0001*) (**Figure 3F**). Taken together, these results suggest that a reduction of PARP trapping may be a common mechanism driving resistance to PARPi.

Expression of *ABCB1 (MDR1), PARG* and *TP53BP1* was assessed using RT-qPCR (**Figure 3G**,**H**). *ABCB1* was undetectable in both PARPi-exposed cells. No significant decrease in *PARG* expression was observed, although a significant increase was observed in the olaparib-exposed cells (*p=0*.*009*), *TP53BP1* expression also remained unchanged.

To assess further whether reduced PARP trapping resulted from lower intracellular drug concentrations, we utilised the intrinsic fluorescence of rucaparib and quantified fluorescence per cell as a proxy for intracellular concentration. There were no significant differences in median population fluorescence values between sensitive and PARPi exposed populations (*p=0*.*9058, p=0*.*9973*) following treatment at 10 μM rucaparib (**Figure 3I,J**).

### PARP trapping is decreased following *in vivo* exposure to olaparib

To investigate the potential changes in PARP trapping *in vivo*, nude mice bearing intraperitoneal OVCAR4 GFP-PARP1-mCherryFP xenografts were treated with olaparib (25 mg/kg) or vehicle for two weeks on a five days on/two days off schedule, (**Figure 4A**). Thereafter, mice were monitored until reaching humane endpoint, at which point tumour cells were isolated from ascitic fluid or peritoneal lavage for subsequent monolayer culturing (**Figure 4B**). There was no difference in retrieved spheroid size in the vehicle and olaparib-treated groups, despite some differences in morphology (**Figure 4C**).

**Figure 4:**
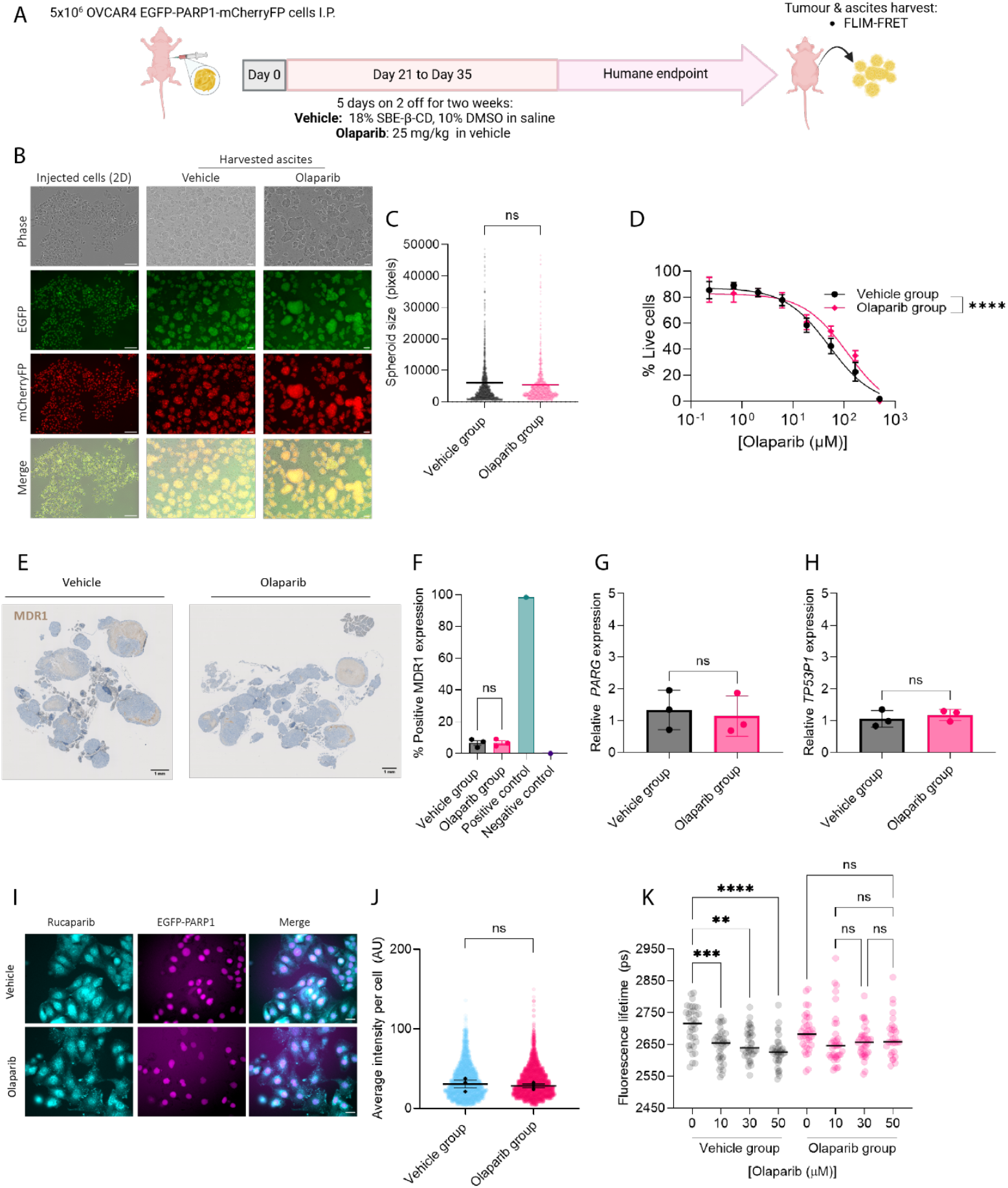
In vivo resistance generation to olaparib leads to a reduction in PARP trapping. **A:** OVCAR4 EGFP-PARP1-mCherryFP cells were injected I.P. into female CD1-nude mice and allowed to grow for 21 days before mice were treated with vehicle or olaparib (5 mg/kg) daily for 14 days. Mice were culled upon reaching humane endpoint and omental tumours and ascites were harvested. **B:** Representative images showing OVCAR4 EGFP-PARP1-mCherryFP cells grown in 2D monolayers before injecting into CD1-nude mice (left) and the ascites harvested from the peritoneal cavity of these mice 90 or 92 days after for vehicle and olaparib treated groups respectively. Images were acquired using the Incucyte S3 with 10X objective. **C:** Quantification of average spheroid area per condition. **D:** Viability assays were performed on the ascites cells grown in 2D, treated with olaparib for 72 h. Survival scores are normalised to vehicle treated controls. Error bars represent SEM, n=3 biological replicates. **E:** Representative IHC images of MDR1 (brown stain) from OVCAR4 EGFP-PARP1-mCherryFP tumours, scale bar = 1mm. **F:** OVCAR4 EGFP-PARP1-mCherryFP omental tumours harvested at day 90 (Vehicle) or 92 (Olaparib) were stained for MDR1 by IHC. The number of MDR1-positive tumour cells was quantified using QuPath^21^. Statistical significance was tested using an unpaired t-test. **G:** RT-qPCR analysis of PARG and **H**: TP53BP1 relative mRNA expression normalised to GAPDH expression. N=3 biological repeats. **I:** Representative images of rucaparib intrinsic fluorescence (left), EGFP-PARP1 (middle) and merged image (right) in the vehicle or olaparib exposed populations, scale bar = 25 μm. **J:** Quantification of median rucaparib signal per cell. N=3 biological repeats. **K:** Fluorescence lifetimes of EGFP measured in FLIM assays of the OVCAR4 EGFP-PARP1-mCherryFP cells that had been harvested from mice ascites of each group. Cells retrieved from vehicle- and olaparib-treated mice were incubated with olaparib (0–50μM) for one hour prior to fluorescence microscopy. Error bars show mean and SEM value, n=3 biological replicates. One-way ANOVA (Kruskal-Wallis) with Dunn’s multiple comparison test was used to assess statistical significance between treatment conditions

Viability assays revealed a significant increase in resistance to olaparib in the olaparib-treated group compared to vehicle-treated (*p*=0.0002) (**Figure 4D**). Expression of MDR1 in OVCAR4 EGFP-PARP1-mCherryFP tumour sections was assessed using IHC (**Figure 4E,F**), showing no significant change between the vehicle (H-score range, 4-9) and olaparib (H-score range, 5-9) treated groups (*p= 0*.*804*). We also assessed expression of *ABCB1, TP53BP1* and *PARG* using RT-qPCR (**Figure 4G,H**). *ABCB1* was again undetectable. Both *TP53BP1* and *PARG* expression remained unchanged between the vehicle and olaparib-exposed groups.

Another possible explanation for observing reduced PARP trapping could be due to reduced intracellular drug concentration due to ECM remodelling or modulation of influx or non-MFR1 efflux pumps. To test this, we again assessed the fluorescence intensity of rucaparib per cell as a readout of intracellular concentration (**Figure 4I,J**) and again found no differences in intracellular concentration, suggesting this is not a mechanism altering PARP trapping in these cells.

Finally, high-content analysis FLIM-FRET assays were performed to evaluate PARP trapping (**Figure 4K**). In vehicle-treated cells, fluorescence lifetimes significantly decreased upon olaparib exposure in a dose-dependent manner, from 2706 ps in untreated cells to 2647, 2652 and 2627 ps at 10, 30 and 50 μM respectively, confirming that dose-dependent PARP trapping remained even after *in vivo* selection and *ex vivo* culturing. By contrast, cells from the olaparib-treated group showed no significant changes in fluorescence lifetime upon olaparib treatment (from 2687 ps in untreated cells to 2664, 2657 and 2675 ps for 10, 30, and 50 μM respectively). These findings demonstrate that the *in vivo* exposure to olaparib not only decreases sensitivity to the drug but also results in reduced levels of PARP trapping.

## Discussion

Here, we describe the development and validation of a novel endogenous PARP1 FRET-based biosensor that enables high-resolution, real-time measurement of PARP trapping in live cells. By combining this biosensor with FLIM readouts, we achieve robust, single-cell quantitative analysis of PARP1 conformational dynamics, serving as a proxy for PARP trapping. This approach can facilitate high-content screening to investigate mechanisms of PARPi resistance. Using this system, we identified reduced PARP trapping in populations with decreased PARPi sensitivity, providing key preclinical evidence linking altered PARP dynamics to resistance. This innovation represents a significant advance, offering new opportunities to study PARPi resistance in clinically relevant settings.

The lack of effective methodologies for studying PARP trapping has limited our understanding of its role in PARPi resistance. Our biosensor directly addresses this gap, offering a tool to measure PARP trapping dynamics in live cells with high-throughput and single-cell resolution. This study demonstrates that the biosensor can discern differences in trapping efficiency across clinically approved PARPi, validating findings from traditional population-based assays such as chromatin immunoprecipitation, while also offering dynamic and single-cell insights. These capabilities are essential for addressing the challenge of overcoming resistance mechanisms in PARP-targeted therapies.

Our findings build upon earlier work by Steffen et al., who developed a fluorescent sensor for PARP1 conformational changes in purified proteins^20^. We extended this method to live cells by employing sequential rounds of CRISPR-Cas9 knock-in for dual-labelling endogenous PARP1 with FRET pairs. Dual-labelling of PARP1 was confirmed using long-read RNA sequencing, where approximately 30% of *PARP1* transcripts covered the full 5386 nucleotide dual-labelled transcript. However, since nanopore sequencing reads unidirectionally (starting from the C-terminus) and the median read length was c.2000 nucleotides, we believe that the results are likely to be a significant underestimation of the true proportion of dual-labelled transcripts and explains the observed enrichment of PARP1-mCherryFP fragments in the long-read sequencing. Based on estimated PARP1 levels (0.2–1 × 10^6^ copies per cell)^22,23^, we estimate there will be least 60,000 copies of the PARP1 FRET biosensor per cell. To our knowledge, this presents one of the first implementations of dual-labelling of a single genomic locus via CRISPR-Cas9. Importantly, we showed clearly that the PARP1 sensor does not impede cellular fitness or PARP1 functionality. Cell size, morphology, growth rate and sensitivity to PARPi and platinum therapy were all unaffected following CRISPR-labelling to express the FRET sensor.

To test the ability to monitor PARP trapping in live cells, we utilised the high-grade serous carcinoma cell line OVCAR4. HCA FLIM-FRET assays enabled us to monitor PARP trapping in thousands of cells per condition with and without drug perturbation. We then found that the sensor could recapitulate previously measured trends in PARP trapping potency of clinically approved PARPi, including veliparib, olaparib, rucaparib and talazoparib. This gave us confidence that we were indeed measuring changes in PARP trapping and facilitated the application of the sensor to assay PARP trapping dynamics of novel disease models where the PARP trapping status was otherwise unknown. This uncovered consistent reductions in PARP trapping measured in populations that had reduced sensitivity to PARPi following continuous exposure to constant dose PARPi for nine weeks *in vitro*.

PARPi-resistant cells exhibited impaired PARP trapping when exposed to both their original treatment and alternative PARP inhibitors, suggesting class-wide adaptations. Additionally, these cells showed decreased cisplatin sensitivity, indicating potential cross-resistance between PARPi and platinum-based therapies.

These findings align with reports of shared resistance mechanisms between PARPi and platinum drugs^24^ and are consistent with clinical observations of reduced platinum efficacy post-PARPi progression in the SOLO2 trial^25^. Further studies are needed to elucidate the molecular pathways driving these resistance mechanisms and their broader implications.

Lastly, we generated biosensor-expressing tumours *in vivo* and measured PARP trapping in cells derived from vehicle or olaparib-treated tumours *ex vivo*, uncovering reductions in PARP trapping in the PARP-treated populations. These results were in accordance with our *in vitro* observations, despite distinct treatment protocols.

Despite its strengths, this study has several limitations. First, our protocols produced only modest IC50 increases in resistance models, in contrast with larger c.10-fold increases observed in prior work where resistance was induced over 11 months using increasing doses of olaparib over time^26^. Thus, it is possible that additional mechanisms, beyond altered PARP trapping, are driven by such treatment protocols which increase dosage over time. However, our protocols were chosen to recreate clinical schedules, where patients receive PARPi at fixed daily doses without any dose escalation. Additionally, OVCAR4 cells are *BRCA1/2* wild-type^27^, and thus cannot reflect all potential resistance mechanisms seen in *BRCA1/2* mutant cells. Our use of OVCAR4 cells was both pragmatic – we could not introduce the sensor into cells with defective HR due to the dependency of our CRISPR KI approach to utilise HR-dependent repair – and clinically-driven – PARPi have activity in HR proficient, *BRCA1/2*-wildtype HGSC both as single agent treatment^28^ and as maintenance^3,4^. Equally, although reversion mutations can be identified in approximately 25% *BRCA*-mutated HGSC progressing on PARPi treatment^8,29^, a significant proportion of resistance in these patients remains unexplained, and unanswered questions remain regarding whether reduced PARP trapping is a direct driver of resistance in BRCA-mutant tumours or other cells with intrinsically higher levels of genomic instability. Thus, we believe that OVCAR4 is a valid cellular model.

Nonetheless, the dynamics of PARP trapping versus BRCA reversion in resistance development warrant further exploration and future efforts will focus on overcoming the restrictions of our CRISPR-knock-in approach.

Several avenues for future research could build upon this work. The biosensor-expressing cells could be grown in spheroids or co-cultures to assess PARP trapping dynamics in *in vitro* models that better recapitulates the tumour microenvironment. Efforts to streamline the CRISPR-Cas9-based dual-labelling process, such as employing a lentiviral system, could facilitate broader adoption of this methodology in varied models, including patient samples and *in vivo*, where experimental tools to measure PARP trapping are currently lacking. Moreover, this would facilitate studies of PARP trapping in *BRCA1/2*-mutated HRD genetic backgrounds. Alternatively, transient expression of *BRCA1/2* during CRISPR transfection may facilitate KI in *BRCA1/2*-mutated cells. The biosensor’s potential extends to benchmarking the trapping potency of novel PARPi and assessing trapping efficiency in clinical samples, paving the way for more personalized therapeutic decisions. Additionally, this platform could support the development of PARPi that exclusively target PARP1, addressing recent findings implicating PARP2 trapping in toxicity^30^.

Ultimately, this biosensor offers a transformative tool for understanding and overcoming PARPi resistance, providing a foundation for novel therapeutic strategies that improve clinical outcomes.

## Acknowledgements

The authors acknowledge benefitting front the technical expertise of Martin Kehoe in establishing the FLIM microscopes. FACS was performed with the aid of the LMS NIHR Flow Cytometry Facility, Imperial College London. Rhodopsin targeting vector (EGFP) was kindly gifted by Professor Izumi Oinuma of the Laboratory of Cell and Molecular Biology, University of Hyogo, Japan. AW69_pDonor^PARP1.TS^ and AM70_pgRNA^PARP1^ plasmids were generated and kindly gifted by Dr Manual A.F.V Gonçalves, Leiden University Medical Center, Department of Cell and Chemical Biology, The Netherlands. *In vivo* experiments were performed by Katie Tyson of the McNeish group. Yuriy Alexandrov of the Physics Department at Imperial College London and the Francis Crick Institute assisted with the implementation and application of the FLIM data analysis tools.

