## Supplementary Figure and Methods for "Assessing PARP trapping dynamics in ovarian cancer using a CRISPR-engineered FRET biosensor"

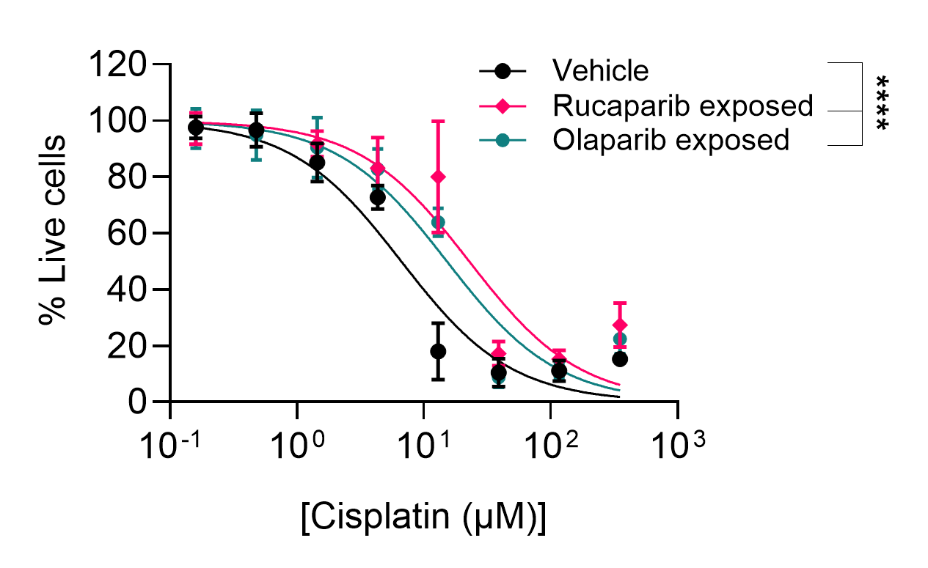

**Supplementary Figure 1: Assessing cisplatin cross-resistance in PARPi treated cells.**

Viability assays were performed on the OVCAR4 EGFP-PARP1-mCherryFP cells following 9 weeks of continuous vehicle, rucaparib or olaparib exposure in vitro. Cells were treated with cisplatin for 72 h. Percentage of live cells are normalised to vehicle treated controls. Error bars represent SEM, n=3 biological replicates.

### Supplementary materials and methods

#### Sulforhodamine B (SRB) cell growth assay

Cell lines were seeded at 7500 cells/well of a 96-well plate. The next day, medium was removed and replaced with drug containing medium, or a vehicle only control (DMSO). 72 h later, cells were fixed with 50% (w/v) trichloroacetic acid (TCA) solution (Sigma-Aldrich T6399; 1 h, 4 °C). Wells were washed 5x with water and dried overnight at room temperature (RT). Cells were stained for 1 h at RT with 0.4% (w/v) SRB (Sigma-Aldrich S1402) solution in 1% (v/v) acetic acid (ThermoFisher 10005920). Plates were washed five times with 1% (v/v) acetic acid to remove excess dye and plates were dried overnight at RT. To solubilise the protein bound SRB dye 100 μl 10 mM Tris Base solution (Sigma-Aldrich T1699) was added to wells and plates were placed on an orbital shaker for 10 min. Absorbance was then measured on a TECAN plate reader at 510 nm. Three technical replicates were used per condition. Media only wells were averaged and subtracted from all wells to remove background. Data were normalised to vehicle control wells.

#### Western Blot

Protein samples were resolved by SDS-PAGE using a Bio-Rad Mini-PROTEAN Tetra System. SDS running buffer (20 mM Tris-HCl, 190 mM glycine, and 0.1% SDS in distilled H₂O) was added to fill the electrophoresis tank, carefully ensuring no leaks between adjacent gels. Samples were loaded, and electrophoresis was conducted at 80 V for 30 minutes, followed by 140 V until the tracking dye reached the gel's bottom.

Protein transfer to a nitrocellulose membrane was performed via wet transfer. Sponges, filter papers, and nitrocellulose membranes were pre-soaked in transfer buffer (20 mM Tris-HCl, 190 mM glycine, 20% methanol in distilled H₂O). Protein transfer was performed at 200 mA for 90 minutes. Ponceau Red (Sigma-Aldrich, P7170) staining was used to confirm protein transfer on the nitrocellulose membrane, followed by three washes in TBS-T to remove excess stain.

Membranes were then blocked in 5% (w/v) BSA (Sigma-Aldrich, A3294) in TBS-T for 1 h at room temperature. Primary antibodies were prepared in BSA solution and incubated with the membranes overnight at 4 °C on a tube roller. After incubation, unbound primary antibodies were removed with three 5-minute washes in TBS-T. Membranes were then incubated with secondary antibodies, diluted in 5% (w/v) powdered milk (Millipore, 70166), for 1 h at room temperature on a tube roller. Excess secondary antibody was removed with three 5-minute washes in TBS-T. Finally, membranes were developed using enhanced chemiluminescence (Cytiva Life Sciences, RPN2106) and imaged with the GE ImageQuant LAS 4000 Biomolecular Imager system.

γH2AX immunofluorescence

Cells were fixed in ice cold methanol for 10 min, permeabilized in permeabilization solution (PS) (0.5% Triton X-100 (Amresco, 0694) in PBS) for 1 hr at RT. Blocking solution (BS) (PS + 10% FBS) was then added for 1 hr at RT. γH2AX-AF488 (Cell Signalling Technology, 20304) staining was performed at 1:10,000 dilution in BS at 4 °C, overnight. SYTO Deep Red (Thermo Fisher, S34900) staining was performed for 1 hr at room temperature (1:1000 in PBS). Three PBS washes were performed, and samples were subsequently imaged using a 100x, 1.4 NA objective Olympus UplanSApo objective lens with HILO illumination. AF488 was imaged with 462 nm laser excitation and SYTO Deep Red was imaged with 635nm laser excitation.

#### IHC MDR1

MDR1/ABCB1 (E1Y7S) 1:300 Ab dilution. (Cell Signalling, 13978).

#### TaqMan Probes:

| **Gene** | **Probe** | **Link** |
| --- | --- | --- |
| *GAPDH* | Hs02786624_g1 | https://www.thermofisher.com/taqman-gene-expression/product/Hs02786624_g1?CID=&ICID=&subtype= |
| *PARG* | Hs00608254_m1 | https://www.thermofisher.com/taqman-gene-expression/product/Hs00608254_m1?CID=&ICID=&subtype= |
| *TP53BP1* | Hs00996827_m1 | https://www.thermofisher.com/taqman-gene-expression/product/Hs00996827_m1?CID=&ICID=&subtype= |
| *ABCB1* | Hs00184500_m1 | https://www.thermofisher.com/taqman-gene-expression/product/Hs00184500_m1?CID=&ICID=&subtype= |

#### Primers used for PCR reactions

**Primer Name// Sequence (5’ -> 3’)**

topA (gRNA) CACC-GTCAATTTTAAGACCTCCCTG

topB (gRNA) CAGTTAAAATTCTGGAGGGAC-CAAA

gfp_f ATGGTGAGCAAGGGCGAGGAGC

gfp_r2 CGGCTACCTCTCCCAAttaccacagTTACTTGTACAGCTCGTCCATGCC

mCherrry_f ATGGTGAGCAAGGGCGAGGAGG

Arm5_F GTGCCACCTGGGCCGGCCATTTAAATggcAGACAAGGATTAGAGGCTG

Arm5_r2 GCTCCTCGCCCTTGCTCACcatGGAGGTCTTAAAATTGAATTTCAGTTTCAGCAG

Arm5_r_mc CCTCCTCGCCCTTGCTCACcatGGAGGTCTTAAAATTGAATTTCAGTTTCAGCAG

arm3_f2 ctgtggtaaTTGGGAGAGGTAGCCG

arm3_r3 GCCGATTCATTAATGCAGCGGCGCcgcTTTCCAAAATCAAAACTAAAGACGAACACTTAG
